# TRPM7 is a critical regulator of pancreatic endocrine development and high-fat diet-induced β-cell proliferation

**DOI:** 10.1101/2020.07.15.204974

**Authors:** Molly K. Altman, Charles M. Schaub, Matthew T. Dickerson, Prasanna K. Dadi, Sarah M. Graff, Thomas J. Galletta, Gautami Amarnath, Ariel S. Thorson, Guoqiang Gu, David A. Jacobson

## Abstract

The melastatin subfamily of the transient receptor potential channels (TRPM) are regulators of pancreatic β-cell function. TRPM7 is the most abundant islet TRPM channel; however, the role of TRPM7 in β-cell function has not been determined. Here, we utilized various spatiotemporal transgenic mouse models to investigate how TRPM7 knockout influences pancreatic endocrine development, proliferation, and function. Ablation of TRPM7 within pancreatic progenitors reduced pancreatic size, as well as α-cell and β-cell mass. This resulted in impaired glucose tolerance due to decreased serum insulin levels. However, ablation of TRPM7 following endocrine specification or in adult mice did not impact endocrine expansion or glucose tolerance. As TRPM7 regulates cell proliferation, we assessed how TRPM7 influences β-cell hyperplasia under insulin resistant conditions. β-cell proliferation induced by high-fat diet was significantly decreased in TRPM7 deficient β-cells. The endocrine roles of TRPM7 may be influenced by cation flux through the channel, and indeed we find that TRPM7 ablation alters β-cell intracellular Mg^2+^. Together, these findings reveal that TRPM7 controls pancreatic progenitor expansion and β-cell proliferation, which is likely due to regulation of Mg^2+^ homeostasis.

**Summary:** This manuscript identifies a critical developmental role for TRPM7 channels in pancreatic progenitor cells. The manuscript also determines that TRPM7 plays a key role in β-cell proliferation under insulin-resistant conditions.

## INTRODUCTION

The melastatin subfamily of the transient receptor potential cation (TRPM) channels are key regulators of β-cell function and apoptosis [1; 2]. For example, Ca^2+^ and Zn^2+^ flux through TRPM2 channels regulates insulin secretion and mitochondrial function, respectively [3; 4]. Activation of TRPM3 by the neuro-steroid hormone pregnenolone-sulfate also enhances insulin biosynthesis and secretion [5]. Moreover, influx of Na^+^ through Ca^2+^-activated TRPM4 and TRPM5 channels enhances glucose-stimulated membrane potential (*V*_m_) depolarization and insulin secretion [6; 7]. TRPM7 is one of the most abundant human β-cell TRPM transcripts; however, its role in islet function is not fully understood [8]. It has been established that TRPM7 channel control of Mg^2+^ homeostasis regulates tissue development [9; 10]. For example, a mutagenesis screen identified TRPM7, along with PDX1 and PTF1, as key determinants of zebrafish pancreatic size [9]. Considering PDX1 and PTF1 play critical roles in pancreatic and β-cell development [11–14], TRPM7 could also serve an important role in the development of the endocrine pancreas.

Magnesium is the most abundant divalent cation in pancreatic β-cells and serves many essential roles, including regulation of the cell cycle [15–19]. TRPM7 activation during G1 phase of the cell cycle corresponds with large increases in Mg^2+^ flux, which stimulates DNA and protein synthesis [20; 21]. However, TRPM7 also modulates Mg^2+^ influx during G0, S, G2, and M phases [20]. Importantly, TRPM7 channel ablation reduces proliferation and leads to cellular senescence, which suggests that Mg^2+^ entry through TRPM7 channels is essential for cell cycle progression [22–24]. Reduced Mg^2+^ influx due to TRPM7 ablation results in increased (e.g. p27^kip1^ and p21^cip1^) or decreased (e.g. cyclin D1, cyclin G1, CDK4) expression of cell cycle regulators [10; 16]. Additionally, loss of TRPM7 Mg^2+^ influx limits phosphorylation-dependent activation of proliferative proteins (e.g. AKT and ERK) [16]. Many of these cell cycle regulators are critical for pancreatic β-cell development and proliferation. For example, p27^Kip1^ ablation increases β-cell mass, while p27^Kip1^ overexpression inhibits β-cell proliferation [25]. Furthermore, activation of AKT and ERK are critical for pregnancy- or high-fat diet (HFD)-induced β-cell proliferation [26–31]. This implicates that TRPM7 control of Mg^2+^ homeostasis may regulate β-cell development and/or proliferation.

Although TRPM7 channels serve a critical role in regulating Mg^2+^ homeostasis, permeability of the channel to Zn^2+^ and Ca^2+^ may also affect β-cell function. Zn^2+^ is critical for insulin crystallization in granules; however, rodent β-cells expressing a loss of function Zn^2+^ transporter (ZnT8) had an increase in insulin secretion, whereas deletion was found to reduce secretion [32; 33]. Since TRPM7 is an outward-rectifying channel and only shows modest Ca^2+^ conductance under physiological *V*_m_, Ca^2+^ influx through TRPM7 channels would be expected to be minimal. Therefore, it is unlikely that Ca^2+^ entry through TRPM7 channels directly affects β-cell Ca^2+^ handling. However, TRPM7 channel control of intracellular Mg^2+^ may indirectly modulate Ca^2+^ handling by modulating voltage-dependent Ca^2+^ (CaV) channel activity [34]. Therefore, it is important to determine how TRPM7 control of β-cell divalent cation flux influences function.

Here we have identified the first roles of TRPM7 channels in pancreatic endocrine cells. The data indicate that TRPM7 channels are key modulators of endocrine development and β-cell proliferation. These studies also determined that TRPM7 limits total islet insulin secretion. These TRPM7 functions are likely mediated by control of Ca^2+^ and Mg^2+^ handling, which are perturbed in TRPM7 deficient β-cells. Therefore, this manuscript reveals that TRPM7 serves as a divalent ion channel in endocrine cells, where it is a critical determinant of pancreatic endocrine development, β-cell proliferation, and insulin secretion.

## RESULTS

### TRPM7 is a critical determinant of pancreatic exocrine and endocrine development

As TRPM7 is essential for zebrafish exocrine development, we investigated if TRPM7 can also affect pancreatic endocrine development. First, TRPM7 was selectively knocked out in pancreatic progenitors utilizing a floxed *TRPM7* exon 17 (TRPM7^fl/fl^,[35]) with mice expressing Pdx1-Cre [36], termed TRPM7KO^Panc^ (Fig. 1A,B and S1). The TRPM7KO^Panc^ mice show significant pancreatic hypoplasia when compared to control mice at embryonic day E16.5 and reduced overall embryonic pancreatic mass (35.6 ± 8.5% reduction, *n* = 4, *P* < 0.05, Fig. 1C). Pancreatic specific TRPM7 ablation also reduced total pancreatic area in adult mice (59.4 ± 20.1%, *n* = 5, P < 0.01, Fig. 1F). Together, these results show that TRPM7 channels are a critical mediator of exocrine development.

**Figure 1.**
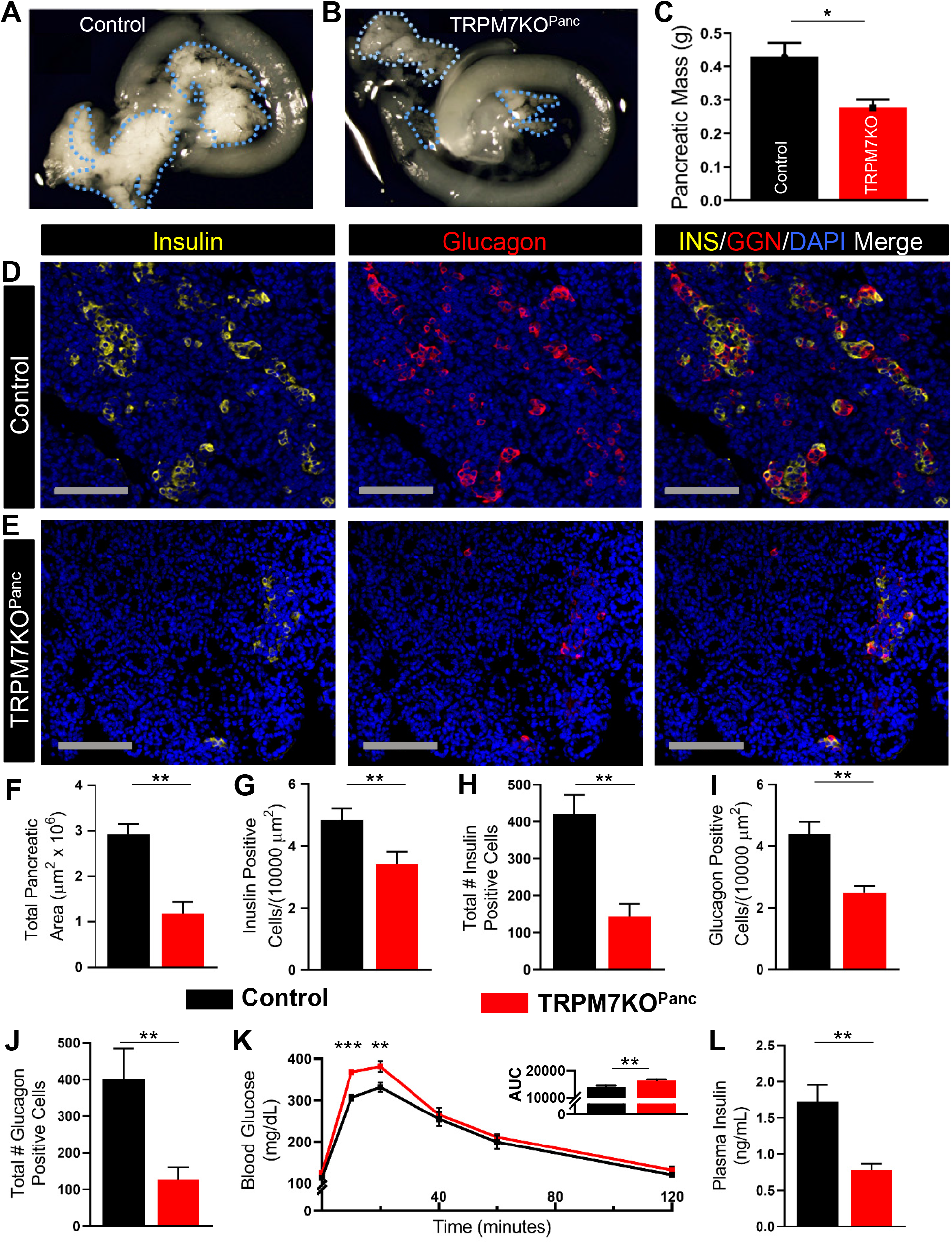
Pancreatic progenitor TRPM7 ablation results in exocrine and endocrine hypoplasia, decreased plasma insulin, and glucose intolerance. Representative images of E16.5 pancreata (dashed blue lines) from control (A) and TRPM7KO^Panc^ (B). (C) Pancreatic weight from control (black) and TRPM7KO^Panc^ (red) mice (*n*=4). Immunofluorescence staining of control (D) and TRPM7KO^Panc^ (E) pancreatic sections for insulin (yellow) and glucagon (red); nuclei (blue), scale bars=100 *u*m. (F) Total pancreatic area (E16.5) of control (*n*=4) vs TRPM7KO^Panc^ (*n*=5). (G) Total number of insulin positive cells per section (control, *n*=7; TRPM7KO^Panc^, *n*=6); (H) Insulin positive cells/1×10^4^ *u*m^2^ (control, *n*=5; TRPM7KO^Panc^, *n*=6). (H) Total number of glucagon positive cells per section (control, *n*=7; TRPM7KO^Panc^, *n*=6); (I) Glucagon positive cells/1×10^4^ *u*m^2^ (control, *n*=4; TRPM7KO^Panc^, *n*=5). (K) Glucose tolerance of TRPM7KO^Panc^ (*n*=7) mice compared to control (*n*=10), inset shows total AUC. (L) Fasting plasma insulin levels in TRPM7KO^Panc^ versus control adult mice. When applicable, data are mean ± SEM and significance is calculated with a two-tailed t-test. *P<0.05; **P<0.01; ***P<0.001.

To better understand the effect of TRPM7 channel ablation on pancreatic endocrine development, we compared the β- and α-cell mass of control and TRPM7KO^Panc^ mice (Fig. 1D,E). In addition to decreased pancreas size, TRPM7KO^Panc^ mice had fewer β-cells (29.7 ± 9.8% decrease; *n* ≥ 5, *P* < 0.05, Fig. 1G) and α-cells (43.4 ± 7.1% decrease, *n* ≥ 4, *P* < 0.01, Fig. 1I) per unit area. Furthermore, there was a reduction in the total number of insulin positive cells (65.9 ± 9.2% decrease, *n* ≥ 6, *P* < 0.01, Fig. 1H) as well as glucagon positive cells (68.6 ± 10.6% decrease, *n* ≥ 6, *P* < 0.05, Fig. 1J) in TRPM7KO^Panc^ pancreatic sections compared to control sections. To assess the impact of reduced endocrine mass on glucose homeostasis, intraperitoneal glucose tolerance tests (IPGTT) were performed. TRPM7KO^Panc^ mice exhibited glucose intolerance compared to control mice (Fig. 1K). Interestingly, fasting plasma insulin was lower in TRPM7KO^Panc^ mice compared to control mice (55.0 ± 8.0% decrease, *n* ≥ 3, *P* < 0.01), which is likely due to reduced β-cell mass and total pancreatic insulin content (Fig. 1L and S2B). While pancreatic TRPM7 channel ablation resulted in impaired glucose tolerance, fasting blood glucose was unaffected. This may indicate that reduced α-cell mass contributes to unaltered fasting glycemia potentially due to reduced fasting glucagon levels. Taken together, these data reveal for the first time that TRPM7 channels are key determinants of pancreatic endocrine development.

### Endocrine progenitor TRPM7 ablation does not alter endocrine mass or glucose homeostasis

To identify the temporal role of TRPM7 channels during pancreatic endocrine development, we next examined if loss of TRPM7 channels in pancreatic endocrine progenitors affects islet development. TRPM7 channels were selectively knocked out in cells following endocrine fate specification by crossing TRPM7^fl/fl^ mice with mice expressing an Ngn3-Cre (TRPM7KO^Endo^, [37; 38]). Interestingly, immunostaining of control and TRPM7KO^Endo^ E16.5 pancreata for insulin and glucagon showed that TRPM7 channel ablation in endocrine progenitors has no effect on total pancreatic area or on β- or α-cell mass (Fig. 2A-G). These data indicate that TRPM7 channels are critical for pancreatic progenitor proliferation, but not differentiation. Furthermore, TRPM7KO^Endo^ mice show equivalent glucose tolerance as controls (Fig. 2H). While this suggests that endocrine specific TRPM7 channel ablation does not directly influence glucose homeostasis, it is possible that TRPM7 channels serve other roles in mature islet cells.

**Figure 2.**
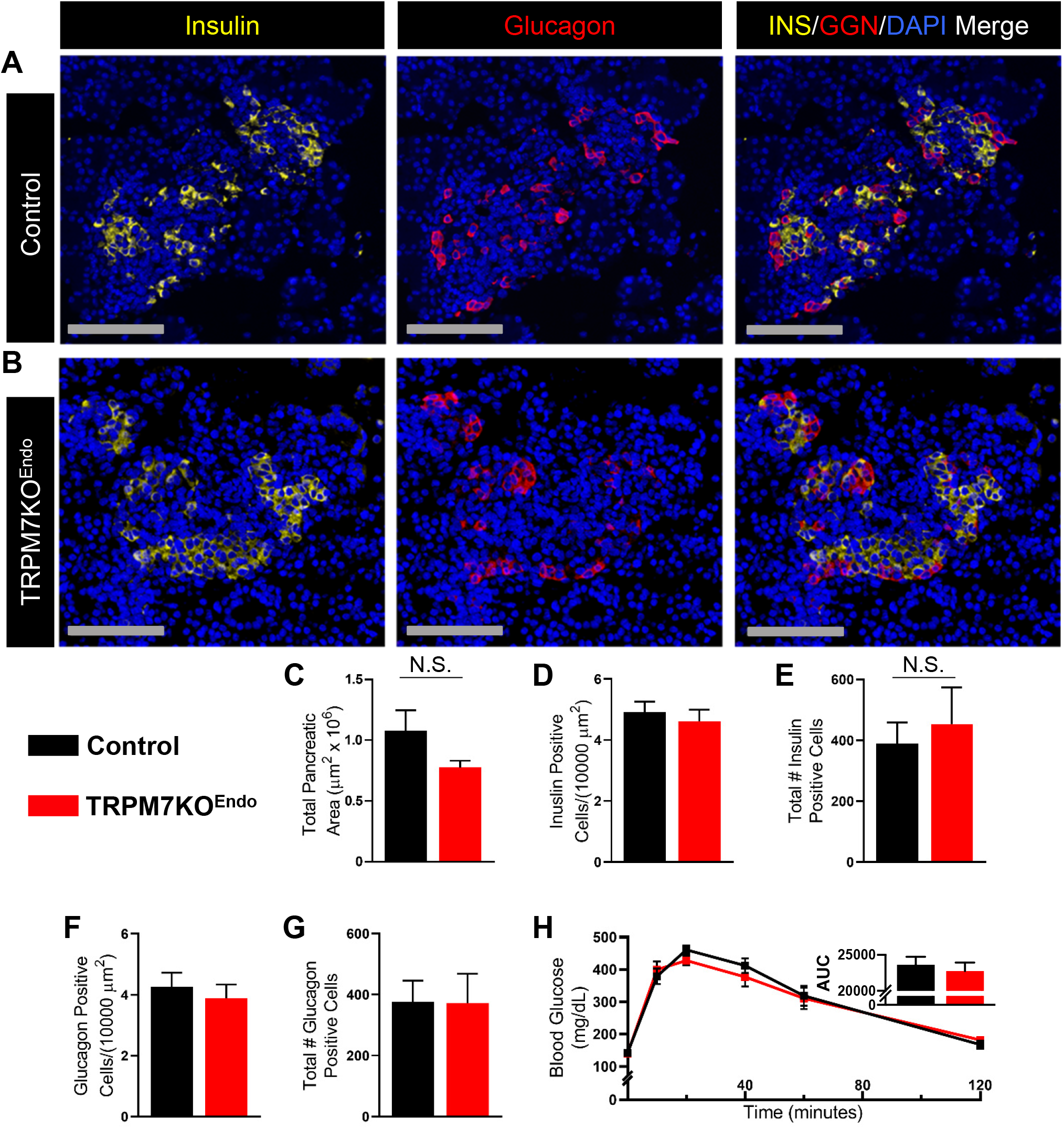
Endocrine progenitor TRPM7 ablation does not impact a- and b-cell mass or glucose tolerance. Immunofluorescence staining of control (A) and TRPM7KO^Endo^ (B) pancreatic sections for insulin (yellow) and glucagon (red); nuclei (blue), scale bars=100 *u*m. (C) Total pancreatic area (E16.5) of control vs TRPM7KO^Endo^ (*n*=3). (D) Total number of insulin positive cells per section (control, *n*=5; TRPM7KO^Endo^, *n*=3); (E) Insulin positive cells/1×10^4^ *u*m^2^ (*n*=3). (F) Total number of glucagon positive cells per section (control, *n*=5; TRPM7KO^Endo^, *n*=3); (G) Glucagon positive cells/1×10^4^ *u*m^2^ (*n*=3). (H) Glucose tolerance of TRPM7KO^Endo^ (*n*=4) mice compared to control (*n*=5), inset shows total AUC. When applicable, data are mean ± SEM and significance is calculated with a two-tailed t-test. N.S., not significant; *P<0.05.

### TRPM7 channels control β-cell insulin secretion

We next investigated the role(s) of TRPM7 channels in mature β-cells. To accomplish this, two β-cell specific conditional KO mouse lines were generated; TRPM7^fl/fl^ mice were crossed with animals expressing MIP-Cre/ERT (TRPM7KO^MIPβ^, [39]) or *Ins1*^*CreERT2*^ (TRPM7KO^INSβ^, [40]). TRPM7 channel ablation specifically in mature β-cells (TRPM7KO^MIPβ^ or TRPM7KO^INSβ^) did not alter glucose tolerance compared to control animals on a chow diet (Fig. 3A,C; Lab Diets, 5LOD); this is also in agreement with the normal glucose tolerance observed in TRPM7KO^Endo^ mice. As TRPM7 is a divalent cation channel that could regulate Ca^2+^ handling, we also assessed TRPM7 channel control of β-cell insulin secretion *ex vivo*. While, β-cell specific TRPM7 channel ablation did not affect insulin secretion at 2- and 7-mM glucose, GSIS (14 mM glucose) was significantly increased (73.7 ± 16.0% increase, *n* = 4, *P* < 0.001, Fig. 3B and S3). TRPM7 channel control of β-cell insulin secretion is also exemplified by a recent report showing that TRPM7 channel knockdown in rat insulinoma cells (INS-1) augments insulin secretion [41]. Taken together, these data suggest that TRPM7 channels not only play an important role in β-cell development, but also in limiting insulin secretion.

**Figure 3.**
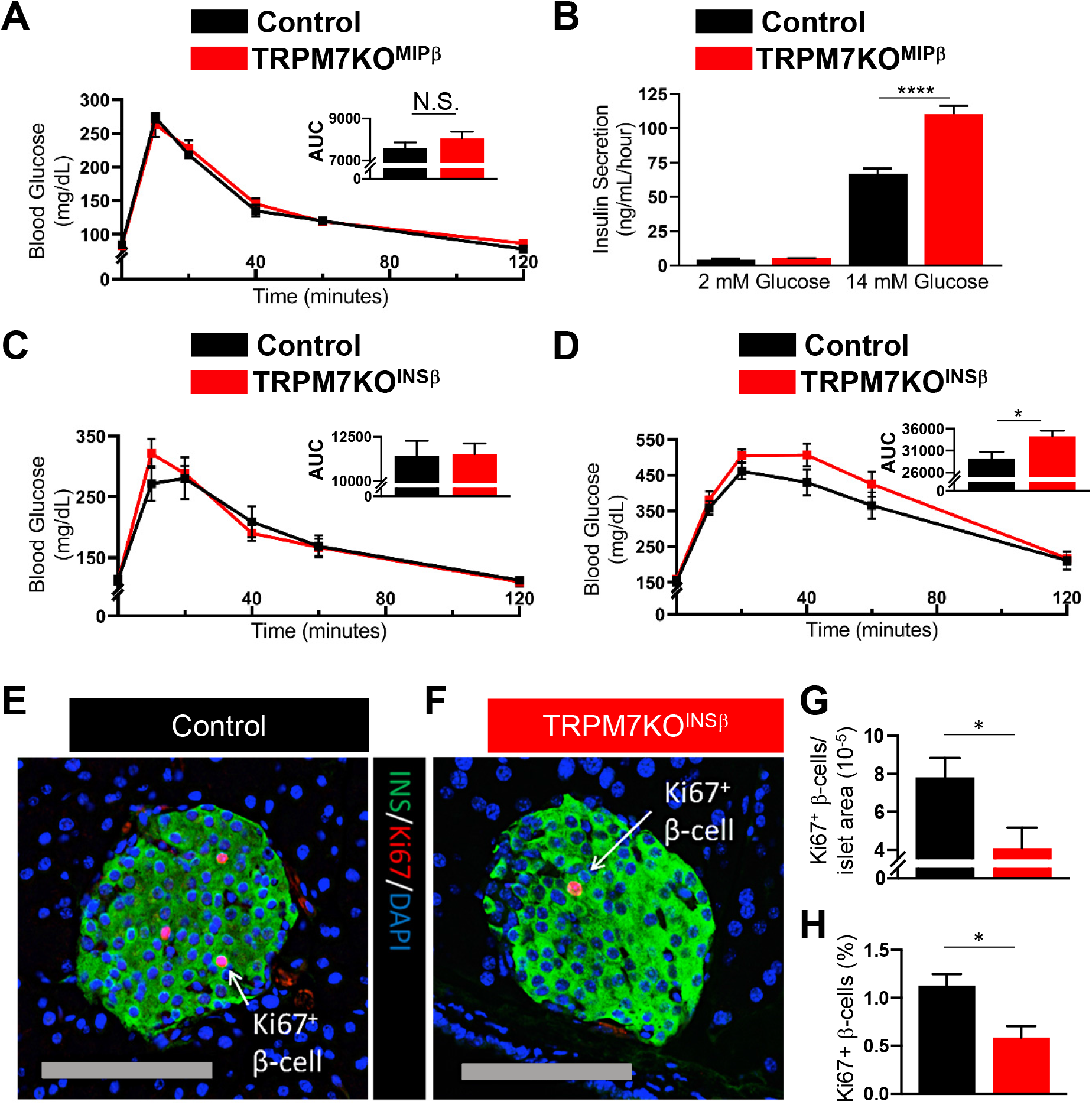
TRPM7 β-cell specific ablation augments GSIS but reduces HFD-induced β-cell proliferation. (A) Glucose tolerance is unaltered in TRPM7KO^MIPβ^ mice (*n*=4) compared to control (*n*=5) (inset shows total AUC), (B) despite enhanced GSIS of isolated islets at 14 mM glucose (*n*=4). (C) TRPM7KO^Insβ^ mice have normal glucose tolerance compared to control (*n*=5), (D) but exposure to HFD (2-weeks) saw increased glucose intolerance (*n*=8); insets show total AUC. (E, F) Immunofluorescent staining of β-cells (green) expressing the Ki-67 proliferation marker (pink) following exposure to HFD; nuclei (blue), scale bars=100 *u*m. (G) Percent of β-cells positive for Ki-67 (*n*=6). (H) Quantification of Ki-67^+^ β-cells/area (*n*=4). When applicable, data are mean ± SEM and significance is calculated with a two-tailed t-test. *P<0.05.; **P<0.01; ***P<0.001.

### TRPM7 channel ablation reduces HFD-induced β-cell proliferation

As our findings show that TRPM7 channels affect embryonic β-cell development, TRPM7 channel control of mature β-cell proliferation was investigated. To accomplish this, adult TRPM7KO^INSβ^ mice were placed on a HFD to promote β-cell proliferation [42]. To confirm insulin resistance following HFD, GTT were first performed. While all animals on the HFD (2-weeks) showed glucose intolerance, TRPM7KO^INSβ^ mice displayed further impairment of glucose tolerance compared to control animals (Fig 3D). We next stained pancreatic sections from TRPM7KO^INSβ^ and control mice on HFD (2 weeks) for Ki67 to determine if changes in β-cell proliferation contribute to glucose intolerance. The number of Ki67+ β-cells per unit area in TRPM7KO^INSβ^ pancreatic sections was significantly decreased compared to control sections (47.7 ± 14.7 decrease, *n* = 4, *P* < 0.05); the percentage of Ki67+ β-cells in TRPM7KO^INSβ^ islets was also reduced relative to control islets (47.9 ± 12.0% decrease, *n* = 6, *P* < 0.05, Fig. 3G,H). Based on these results, TRPM7 channel ablation in mature β-cells does not affect glucose homeostasis under physiological conditions. Whereas, during conditions of metabolic stress (HFD), β-cell TRPM7 KO leads to glucose intolerance, which is due in part to reduced β-cell proliferation. Additionally, it is also possible that enhanced insulin secretion observed in TRPM7KO^INSβ^ animals exacerbates glucose intolerance in response to a HFD [43; 44].

### TRPM7 forms functional channels that control β-cell Ca^2+^ and Mg^2+^ handing

Because TRPM7 has been shown to affect cellular function, development, and proliferation by controlling divalent cation homeostasis, we next examined if TRPM7 channels are functional on the β-cell plasma membrane [18; 35; 45]. Total β-cell TRPM7 currents were monitored in response to a voltage ramp from −120 mV to +60 mV in either the presence (black circles) or absence (grey circles) of extracellular divalent cations (Fig. 4A). This was then repeated for TRPM7KO^Panc^ β-cells under the same conditions (Fig. 4B). The current that was activated under monovalent conditions was considered the TRPM7 current (reference [46]). This was supported by a significant reduction in the TRPM7 current under hyperpolarized (−120 to −65 mV) and depolarized (+25 to +60 mV) conditions in TRPM7KO^Panc^ β-cells compared to controls (Fig. 4C; P<0.05). These data indicate that TRPM7 could contribute to divalent flux across the β-cell plasma membrane.

**Figure 4:**
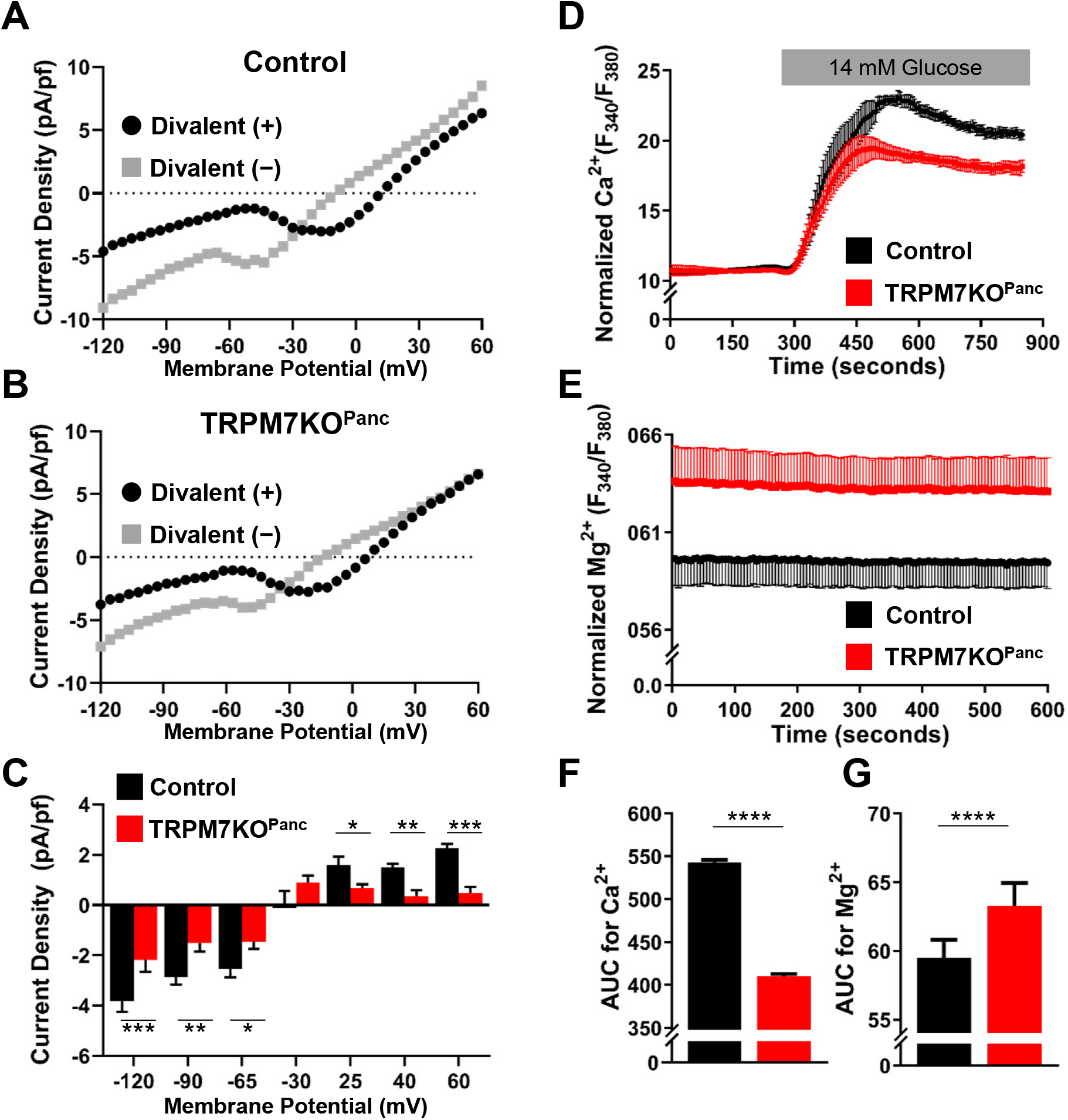
TRPM7 enhances β-cell Mg^2+^ efflux and limits Ca^2+^ entry. Representative β-cell TRPM7 current activation in response to divalent cation removal from (A) control and (B) TRPM7KO^Panc^ β-cells. (C) Divalent activated cation current from control (black, *n*=8) and TRPM7KO^Panc^ (red, *n*=7) β-cells at the specified membrane potentials. (D) Relative glucose-stimulated (11 mM) cytosolic Ca^2+^ responses in control and TRPM7KO^Panc^ islets using Fura-2 (*n*=4). (E) Relative glucose-stimulated (11 mM) cytosolic Ca^2+^ responses in control and TRPM7KO^Panc^ islets using Mag-Fura-2 (*n*=4). (F, G) Area under the curve quantification of cytosolic Ca^2+^ (500-800 seconds) and Mg^2+^ (0-600 seconds) of control (black) and TRPM7KO^Panc^ (red) β-cells/islets. When applicable, data are mean ± SEM and significance is calculated with a two-tailed t-test. *P<0.05.; **P<0.01; ***P<0.001.

As β-cell Ca^2+^ influx is required for GSIS, TRPM7 channel-mediated regulation of β-cell Ca^2+^ handling was assessed. Glucose-stimulated Ca^2+^ influx into TRPM7KO^Panc^ islets was decreased following treatment with high (14 mM) glucose compared to control islets (24.1 ± 0.6% decrease, *n* = 4, P<0.001, Fig. 4E,G). To determine if TRPM7 channel-mediated enhancement of β-cell intracellular Ca^2+^ concentration ([Ca^2+^]_i_) was due to its impact on β-cells during development, glucose-stimulated Ca^2+^ influx into TRPM7KO^MIPβ^ islets was also measured. Similar to total pancreatic TRPM7 channel ablation, glucose-stimulated Ca^2+^ influx was reduced in TRPM7KO^MIPβ^ islets compared to control islets (Fig. S4A,B). TRPM7 channels are also highly permeable to Zn^2+^ ions, which have been shown to regulate GSIS. However, β-cell glucose-stimulated Zn^2+^ influx was identical in TRPM7KO^MIPβ^ and control islets (Fig. S4D). Together, these data indicate that TRPM7 channels augment glucose-stimulated β-cell Ca^2+^ handling, which suggests that TRPM7 mediated reductions in GSIS are not due to an increase in β-cell [Ca^2+^]_i_.

Magnesium plays a critical role in the cell cycle as well as signaling associated with insulin secretion, thus, TRPM7 channel regulation of islet Mg^2+^ homeostasis was next assessed. Cytosolic Mg^2+^ was elevated in TRPM7KO^Panc^ islets compared to control islets following removal of extracellular Mg^2+^ (6.3 ± 0.1 % increase, *n* = 4, P<0.05, Fig. 4D,F). Cytosolic Mg^2+^ was also elevated when TRPM7 channels were ablated in mature β-cells (TRPM7KO^MIPβ^, Fig. S4C). This suggests that under these conditions TRPM7 either directly regulates Mg^2+^ efflux or indirectly reduces β-cell intracellular Mg^2+^ concentration ([Mg^2+^]_i_). For example, Mg^2+^ efflux is reduced during GSIS, which is further enhanced in media lacking Mg^2+^ [47]; therefore, enhanced GSIS in TRPM7 KO islets could limit Mg^2+^ efflux. Magnesium is a positive regulator of adenylate cyclase activity, and thus, the elevated Mg^2+^ could raise cAMP levels in TRPM7 KO β-cells resulting in amplification of GSIS [48–50]. It is also well-established that TRPM7 activity is dependent on intracellular Mg^2+^. As intracellular Mg^2+^ drops in type-2 diabetes (T2D) due to insulin resistance [51; 52], reduced Mg^2+^ dependent changes in TRPM7 activity could be responsible, in part, for the reduction in β-cell proliferation that occurs with prolonged HFD or increased duration of T2D [53].

TRPM7 control of Mg^2+^ homeostasis likely limits GSIS and may also influence β-cell proliferation. For example, Mg^2+^ influx through TRPM7 channels activates phosphoinositide 3-kinase dependent signaling and AKT phosphorylation, which has been shown to play a critical role in β-cell proliferation [54; 55]. AKT phosphorylation is part of insulin signaling cascade and KO of β-cell insulin receptor abolishes β-cell hyperplasia in response to HFD [56]. Similarly, KO of β-cell AKT1 prevents β-cell hyperplasia in response to HFD [54]. Moreover, expression of a constitutively active AKT1 significantly increases β-cell proliferation leading to increased β-cell mass [28; 57]. Therefore, reduced AKT phosphorylation in TRPM7 deficient β-cells may limit proliferation in response to HFD. Additionally, perturbed Mg^2+^ homeostasis in TRPM7 deficient β-cells may impact other cell cycle pathways controlled by Mg^2+^ such as CDK2 kinase activity, expression of kinase inhibitor genes (e.g. p21^cip1^ and p27^kip1^), DNA synthesis, and/or cytoskeletal remodeling [58–61]. Finally, it has been shown that lymphocytes without TRPM7 channels enter a quiescent state, potentially due to a large reduction in aerobic glycolysis necessary for sustained proliferation. Considering Mg^2+^ is required for sequential enzymatic reactions within the glycolytic pathway, this may account for the reduced proliferation in the TRPM7KO^Ins^ under stressful conditions [17; 62–64].

TRPM7 control of Mg^2+^ plays a critical role in zebrafish pancreatic development and may also impact endocrine development. Indeed, the role of TRPM7 on endocrine cell development during E8.5-12.5 is striking, as TRPM7 KO resulted a greater than 60% reduction α- and β-cell mass (Fig. 1H,J). During early pancreatic development there is a large expansion of the multipotent progenitor cells, which are a critical determinant of adult pancreatic size [65]. Thus, it appears TRPM7 regulates the cell cycle of pancreatic progenitors, presumably through Mg^2+^ homeostasis. Conversely, there are no observed deleterious effects on endocrine mass or glucose tolerance following loss of TRPM7 during endocrine progenitor specification. Therefore, Mg^2+^ is likely not a critical factor in endocrine commitment and exit from the cell cycle [66; 67]. This temporal developmental effect of TRPM7 is supported by studies on neural, kidney and pigment cells, where TRPM7 is critical for early organ development during embryogenesis but has a minimal effect on terminal differentiation [68]. However, TRPM7 may also influence pancreatic development through maintenance of a pluripotent stem cell population by controlling expression of OCT4, NANOG, and STAT3 similarly to thymocytes) [35]. Taken together, our data indicate TRPM7 plays an essential role in early pancreatic development as well as HFD-induced β-cell proliferation. Therefore, these data illustrate that TRPM7 could be a useful target for increasing β-cell mass and/or enhancing islet cell production from induced pluripotent stem cells.

## METHODS

### Mouse Pancreatic and β-cell specific TRPM7 ablation

For spatiotemporal TRPM7 ablation, transgenic animals were utilized by crossing 129S4/SvJae mice containing LoxP sites inserted around exon 17 in the *TRPM7* gene (TRPM7^fl/fl^, STOCK Trpm7tm1Clph/J, The Jackson Laboratory, 018784) [35] with various Cre-recombinase lines. For pancreatic progenitor specific ablation, TRPM7^fl/fl^ were crossed with C57BL/6 mice having a Cre-recombinase under the Pdx1 promoter (B6.FVB-Tg(Pdx1-cre)6Tuv/J) [35]. Endocrine specific ablation mice were created by crossing TRPM7^fl/fl^ with Ngn3-cre mice [37]. β-cell-specific TRPM7 ablation are a cross between TRPM7^fl/fl^ and C57BL/6 mice with a tamoxifen activated CreERT-recombinase expressed in β-cells via an insulin promoter (MIP-CreERT, B6.Cg-Tg(Ins1-cre/ERT)1Lphi/J, the Jackson Laboratory, 024709; Ins1^CreERT2^, B6(Cg)-*Ins1*^*tm2.1(cre/ERT2)Thor*^/J, the Jackson Laboratory, 026802) [39; 40]. To induce translocation of the CreERT into the nucleus, CreERT mice were treated with tamoxifen (2 mg·ml^−1^; RPI, Mount Prospect, IL) every other day 5 times as indicated. Controls were TRPM7^fl/fl^ animals, MIP-CreERT or Ins1^CreERT2^ mice; all were treated with tamoxifen identically to the TRPM7^fl/fl^-MIP-CreERT and TRPM7^fl/fl^-*Ins1*^*CreERT2*^ mice.

### Mouse Diets and Glucose Tolerance Testing

Mice were placed either on a normal chow diet or a high fat diet (HFD, 60 kcal% fat, D12492 Research Diets, Inc.) and monitored for glucose tolerance. The glucose tolerance test (GTT) was performed as described previously by injecting 2 mg/kg dextrose (animals on a normal chow diet) or 2 mg/kg (animals on a HFD) and monitoring blood glucose at the indicated time points post glucose injection after an approximately 5-6 hour fast [69]. GTT was performed between 12-19 weeks of age. A cohort of mice was also placed on HFD at 15 weeks of age for 2 weeks; at this time, which another GTT was performed.

### Mouse Islet and β-Cell Isolation

Mouse islets were isolated by digesting the pancreas with collagenase P (Roche) and performing density gradient centrifugation as previously described [70] . Islets were plated or dissociated in 0.005% trypsin, placed on glass coverslips, and cultured for 16 h in RPMI 1640 medium supplemented with 10% fetal calf serum, concentrations of glucose-specified, 100 international units ml^−1^ penicillin, and 100 mg ml^−1^ streptomycin. Dissociated β-cells were specifically used in all voltage clamp experiments recording Mg^2+^ currents. β-Cells on the periphery of intact islets were recorded in current clamp mode in all the membrane potential recordings. Cells and islets were maintained in a humidified incubator at 37 °C under an atmosphere of 95% air and 5% CO_2_. Islet cells were incubated in 2 mM glucose with Fura 2AM (Molecular Probes) for 25 min. before Ca^2+^ imaging.

### Whole Cell Voltage Clamp Electrophysiological Recordings

TRPM7KO^Panc^ islets were dispersed into single cells and cultured overnight at 37°C, 5% CO_2_. Recording pipettes (8-10 MΩ) were backfilled with intracellular solution containing (mM): 140.0 CsCl, 1.0 MgCl_2_, 10.0 EGTA, and 10.0 HEPES (pH 7.2, adjusted with CsOH) supplemented with 3.7 Mg-ATP. Cells were patched in extracellular buffer containing (mM) 119.0 NaCl, 4.7 KCl, 25.0 HEPES, 1.2 MgSO_4_, 1.2 KH_2_PO_4_, 2.0 CaCl_2_, (pH 7.4, adjusted with NaOH) supplemented with 14.0 mM glucose. After a whole-cell configuration was established (seal resistance > 1 GΩ) the bath solution was exchanged (3 minutes) for divalent cation-free buffer containing (mM) 119.0 NaCl, 4.7 KCl, 25.0 HEPES, 1.2 KH_2_PO_4_, and 10.0 tetraethylammonium chloride (TEA) supplemented with 14.0 mM glucose (pH 7.4, adjusted with NaOH). A voltage-clamp protocol was employed to record whole-cell currents with and without extracellular divalent cations in response to a voltage ramp from −120 to 60 mV (1 second interval) using an Axopatch 200B amplifier with pCLAMP10 software (Molecular Devices) as previously described [71; 72]. TRPM7 currents were calculated by subtracting currents without divalents from currents with divalents. Data were analyzed with Clampfit 10 (Molecular Devices) and Microsoft Excel software.

### Measurement of Cytoplasmic Calcium

Islet cell clusters were incubated (25 min at 37 °C) in RPMI supplemented with 2 μM Fura-2 acetoxymethyl ester (Molecular Probes, Eugene, OR). Fluorescence imaging was performed using a Nikon Eclipse TE2000-U microscope equipped with an epifluorescence illuminator (SUTTER, Inc.) a CCD camera (HQ2, Photometrics, Inc.) and Nikon Elements software (NIKON, Inc.). Cells were perifused at 37 °C at a flow of 2 ml/min with appropriate KRB-based solutions that contained glucose concentrations and compounds specified in the figures. Relative Ca^2+^ concentrations were quantified every 5 s by determining the ratio of emitted fluorescence intensities at excitation wavelengths of 340 and 380 nm (F_340_/F_380_). The relative Ca^2+^ is plotted as the average FURA-2 ratio (F_340_/F_380_) of each experimental group ± the S.E. The averaged area under the curve (AUC) measurement for FURA-2 ratios during the indicated time period (in minutes) was plotted as a bar graph. Data were analyzed using Excel and GraphPad Prism software and compared by Student’s *t* test.

### Measurement of Cytoplasmic Magnesium

Islet cell clusters were incubated (35 min at 37 °C) in RPMI supplemented with Mag-fura-2 AM an intracellular magnesium indicator that is ratiometric and UV light-excitable. This acetoxymethyl (AM) ester form is useful for noninvasive intracellular loading (Invitrogen M1292). Islet clusters were washed 3 times with 0 Mg^2+^ KRB-based solution. The 3^rd^ wash remained on the cells for the approximately 25-30-minute preincubation prior to imaging. Fluorescence imaging and analysis of data was performed on a Nikon Eclipse scope exactly as described for our cytoplasmic calcium measurements.

### Plasma Insulin Measurements

Animals were starved for approximately 4 hours prior to glucose injections to stimulate insulin production and then blood was collected for Insulin measurements. Blood was collected in EDTA-Heprin coated VAT tubes (Sarstedt, Germany). VAT Tubes were spun down with centrifugation at 3000 RPM for 10 minutes at 4 °C and serum supernatant was collected into fresh Eppendorf tubes. Serum samples were assayed by ELISA processed by the Vanderbilt Hormone Core.

### Islet Immunofluorescence Staining

Pancreas cubes from TRPM7^fl/fl^-*Ins1*^*CreERT2*^ animals were fixed in 4% paraformaldehyde, paraffin embedded and cut into 5-μm sections on a microtome. Islet β-cells were stained using insulin (Millipore), glucagon (Sigma), and Ki-67 (Abcam) antibodies at 1:300 in combination with fluorescent isothiocyanate-conjugated secondary antibodies at 1:300 (Jackson ImmunoResearch Laboratories) together with DAPI nuclear stain (Invitrogen).

### Statistical Analysis

Data was analyzed using Excel and GraphPad Prism software and compared by student t-test and multiple ANOVA. Statistical analysis was performed by using either the two-way ANOVA followed by the Bonferroni Test or the Student’s T-Test. A P-value of <0.05 was considered statistically significant.

## REFERENCES

1. Uchida, K. and Tominaga, M. (2011). The role of thermosensitive TRP (transient receptor potential) channels in insulin secretion. Endocr J 58, 1021–1028.

2. Islam, M. S. (2020). Molecular Regulations and Functions of the Transient Receptor Potential Channels of the Islets of Langerhans and Insulinoma Cells. Cells 9.

3. Li, X., Yang, W. and Jiang, L. H. (2017). Alteration in Intracellular Zn(2+) Homeostasis as a Result of TRPM2 Channel Activation Contributes to ROS-Induced Hippocampal Neuronal Death. Front Mol Neurosci 10, 414.

4. Li, F., Munsey, T. S. and Sivaprasadarao, A. (2017). TRPM2-mediated rise in mitochondrial Zn(2+) promotes palmitate-induced mitochondrial fission and pancreatic beta-cell death in rodents. Cell Death Differ 24, 1999–2012.

5. Thiel, G., Muller, I. and Rossler, O. G. (2013). Signal transduction via TRPM3 channels in pancreatic beta-cells. J Mol Endocrinol 50, R75–83.

6. Cheng, H., Beck, A., Launay, P., Gross, S. A., Stokes, A. J., Kinet, J. P., Fleig, A. and Penner, R. (2007). TRPM4 controls insulin secretion in pancreatic beta-cells. Cell Calcium 41, 51–61.

7. Brixel, L. R., Monteilh-Zoller, M. K., Ingenbrandt, C. S., Fleig, A., Penner, R., Enklaar, T., Zabel, B. U. and Prawitt, D. (2010). TRPM5 regulates glucose-stimulated insulin secretion. Pflugers Arch 460, 69–76.

8. Marabita, F. and Islam, M. S. (2017). Expression of Transient Receptor Potential Channels in the Purified Human Pancreatic beta-Cells. Pancreas 46, 97–101.

9. Yee, N. S., Lorent, K. and Pack, M. (2005). Exocrine pancreas development in zebrafish. Dev Biol 284, 84–101.

10. Yee, N. S., Zhou, W. and Liang, I. C. (2011). Transient receptor potential ion channel Trpm7 regulates exocrine pancreatic epithelial proliferation by Mg2+-sensitive Socs3a signaling in development and cancer. Dis Model Mech 4, 240–254.

11. Offield, M. F., Jetton, T. L., Labosky, P. A., Ray, M., Stein, R. W., Magnuson, M. A., Hogan, B. L. and Wright, C. V. (1996). PDX-1 is required for pancreatic outgrowth and differentiation of the rostral duodenum. Development 122, 983–995.

12. Krapp, A., Knofler, M., Frutiger, S., Hughes, G. J., Hagenbuchle, O. and Wellauer, P. K. (1996). The p48 DNA-binding subunit of transcription factor PTF1 is a new exocrine pancreas-specific basic helix-loop-helix protein. EMBO J 15, 4317–4329.

13. Kawaguchi, Y., Cooper, B., Gannon, M., Ray, M., MacDonald, R. J. and Wright, C. V. (2002). The role of the transcriptional regulator Ptf1a in converting intestinal to pancreatic progenitors. Nat Genet 32, 128–134.

14. Burlison, J. S., Long, Q., Fujitani, Y., Wright, C. V. and Magnuson, M. A. (2008). Pdx-1 and Ptf1a concurrently determine fate specification of pancreatic multipotent progenitor cells. Dev Biol 316, 74–86.

15. Sgambato, A., Wolf, F. I., Faraglia, B. and Cittadini, A. (1999). Magnesium depletion causes growth inhibition, reduced expression of cyclin D1, and increased expression of P27Kip1 in normal but not in transformed mammary epithelial cells. J Cell Physiol 180, 245–254.

16. Fang, L., Zhan, S., Huang, C., Cheng, X., Lv, X., Si, H. and Li, J. (2013). TRPM7 channel regulates PDGF-BB-induced proliferation of hepatic stellate cells via PI3K and ERK pathways. Toxicol Appl Pharmacol 272, 713–725.

17. Sahni, J. and Scharenberg, A. M. (2008). TRPM7 ion channels are required for sustained phosphoinositide 3-kinase signaling in lymphocytes. Cell Metab 8, 84–93.

18. Sahni, J., Tamura, R., Sweet, I. R. and Scharenberg, A. M. (2010). TRPM7 regulates quiescent/proliferative metabolic transitions in lymphocytes. Cell Cycle 9, 3565–3574.

19. Yu, Y., Chen, S., Xiao, C., Jia, Y., Guo, J., Jiang, J. and Liu, P. (2014). TRPM7 is involved in angiotensin II induced cardiac fibrosis development by mediating calcium and magnesium influx. Cell Calcium 55, 252–260.

20. Tani, D., Monteilh-Zoller, M. K., Fleig, A. and Penner, R. (2007). Cell cycle-dependent regulation of store-operated I(CRAC) and Mg2+-nucleotide-regulated MagNuM (TRPM7) currents. Cell Calcium 41, 249–260.

21. Rubin, H. (2005). The membrane, magnesium, mitosis (MMM) model of cell proliferation control. Magnes Res 18, 268–274.

22. Yee, N. S., Zhou, W., Lee, M. and Yee, R. K. (2012). Targeted silencing of TRPM7 ion channel induces replicative senescence and produces enhanced cytotoxicity with gemcitabine in pancreatic adenocarcinoma. Cancer Lett 318, 99–105.

23. Brandao, K., Deason-Towne, F., Perraud, A. L. and Schmitz, C. (2013). The role of Mg2+ in immune cells. Immunol Res 55, 261–269.

24. Beesetty, P., Wieczerzak, K. B., Gibson, J. N., Kaitsuka, T., Luu, C. T., Matsushita, M. and Kozak, J. A. (2018). Inactivation of TRPM7 kinase in mice results in enlarged spleens, reduced T-cell proliferation and diminished store-operated calcium entry. Sci Rep 8, 3023.

25. Rachdi, L., Balcazar, N., Elghazi, L., Barker, D. J., Krits, I., Kiyokawa, H. and Bernal-Mizrachi, E. (2006). Differential effects of p27 in regulation of beta-cell mass during development, neonatal period, and adult life. Diabetes 55, 3520–3528.

26. Gupta, R. K., Gao, N., Gorski, R. K., White, P., Hardy, O. T., Rafiq, K., Brestelli, J. E., Chen, G., Stoeckert, C. J., Jr. and Kaestner, K. H. (2007). Expansion of adult beta-cell mass in response to increased metabolic demand is dependent on HNF-4alpha. Genes & development 21, 756–769.

27. Bernal-Mizrachi, E., Fatrai, S., Johnson, J. D., Ohsugi, M., Otani, K., Han, Z., Polonsky, K. S. and Permutt, M. A. (2004). Defective insulin secretion and increased susceptibility to experimental diabetes are induced by reduced Akt activity in pancreatic islet beta cells. J Clin Invest 114, 928–936.

28. Bernal-Mizrachi, E., Wen, W., Stahlhut, S., Welling, C. M. and Permutt, M. A. (2001).Islet beta cell expression of constitutively active Akt1/PKB alpha induces striking hypertrophy, hyperplasia, and hyperinsulinemia. J Clin Invest 108, 1631–1638.

29. Chen, H., Gu, X., Liu, Y., Wang, J., Wirt, S. E., Bottino, R., Schorle, H., Sage, J. and Kim, S. K. (2011). PDGF signalling controls age-dependent proliferation in pancreatic beta-cells. Nature 478, 349–355.

30. Balcazar, N., Sathyamurthy, A., Elghazi, L., Gould, A., Weiss, A., Shiojima, I., Walsh, K. and Bernal-Mizrachi, E. (2009). mTORC1 activation regulates beta-cell mass and proliferation by modulation of cyclin D2 synthesis and stability. J Biol Chem 284, 7832–7842.

31. Alejandro, E. U., Gregg, B., Wallen, T., Kumusoglu, D., Meister, D., Chen, A., Merrins, M. J., Satin, L. S., Liu, M., Arvan, P., et al. (2014). Maternal diet-induced microRNAs and mTOR underlie beta cell dysfunction in offspring. J Clin Invest.

32. Kleiner, S., Gomez, D., Megra, B., Na, E., Bhavsar, R., Cavino, K., Xin, Y., Rojas, J., Dominguez-Gutierrez, G., Zambrowicz, B., et al. (2018). Mice harboring the human SLC30A8 R138X loss-of-function mutation have increased insulin secretory capacity. Proc Natl Acad Sci U S A 115, E7642–E7649.

33. Hardy, A. B., Wijesekara, N., Genkin, I., Prentice, K. J., Bhattacharjee, A., Kong, D., Chimienti, F. and Wheeler, M. B. (2012). Effects of high-fat diet feeding on Znt8-null mice: differences between beta-cell and global knockout of Znt8. Am J Physiol Endocrinol Metab 302, E1084–1096.

34. Zierler, S., Hampe, S. and Nadolni, W. (2017). TRPM channels as potential therapeutic targets against pro-inflammatory diseases. Cell Calcium 67, 105–115.

35. Jin, J., Desai, B. N., Navarro, B., Donovan, A., Andrews, N. C. and Clapham, D. E. (2008). Deletion of Trpm7 disrupts embryonic development and thymopoiesis without altering Mg2+ homeostasis. Science 322, 756–760.

36. Hingorani, S. R., Petricoin, E. F., Maitra, A., Rajapakse, V., King, C., Jacobetz, M. A., Ross, S., Conrads, T. P., Veenstra, T. D., Hitt, B. A., et al. (2003). Preinvasive and invasive ductal pancreatic cancer and its early detection in the mouse. Cancer Cell 4, 437–450.

37. Gu, G., Dubauskaite, J. and Melton, D. A. (2002). Direct evidence for the pancreatic lineage: NGN3+ cells are islet progenitors and are distinct from duct progenitors. Development 129, 2447–2457.

38. Mastracci, T. L., Anderson, K. R., Papizan, J. B. and Sussel, L. (2013). Regulation of Neurod1 contributes to the lineage potential of Neurogenin3+ endocrine precursor cells in the pancreas. PLoS Genet 9, e1003278.

39. Wicksteed, B., Brissova, M., Yan, W., Opland, D. M., Plank, J. L., Reinert, R. B., Dickson, L. M., Tamarina, N. A., Philipson, L. H., Shostak, A., et al. (2010). Conditional gene targeting in mouse pancreatic ss-Cells: analysis of ectopic Cre transgene expression in the brain. Diabetes 59, 3090–3098.

40. Thorens, B., Tarussio, D., Maestro, M. A., Rovira, M., Heikkila, E. and Ferrer, J. (2015). Ins1(Cre) knock-in mice for beta cell-specific gene recombination. Diabetologia 58, 558–565.

41. Gommers, L. M. M., Hill, T. G., Ashcroft, F. M. and de Baaij, J. H. F. (2019). Low extracellular magnesium does not impair glucose-stimulated insulin secretion. PLoS One 14, e0217925.

42. Golson, M., Misfeldt, A. A., Kopsombut, U., Petersen, C. and Gannon, M. (2010). High fat diet regulation of β-cell proliferation and β-cell mass. The open endocrinology journal 4.

43. Lillioja, S., Mott, D. M., Howard, B. V., Bennett, P. H., Yki-Jarvinen, H., Freymond, D., Nyomba, B. L., Zurlo, F., Swinburn, B. and Bogardus, C. (1988). Impaired glucose tolerance as a disorder of insulin action. Longitudinal and cross-sectional studies in Pima Indians. N Engl J Med 318, 1217–1225.

44. Warram, J. H., Martin, B. C., Krolewski, A. S., Soeldner, J. S. and Kahn, C. R. (1990). Slow glucose removal rate and hyperinsulinemia precede the development of type II diabetes in the offspring of diabetic parents. Ann Intern Med 113, 909–915.

45. Ryazanova, L. V., Rondon, L. J., Zierler, S., Hu, Z., Galli, J., Yamaguchi, T. P., Mazur, A., Fleig, A. and Ryazanov, A. G. (2010). TRPM7 is essential for Mg(2+) homeostasis in mammals. Nat Commun 1, 109.

46. Zhang, Y. H., Sun, H. Y., Chen, K. H., Du, X. L., Liu, B., Cheng, L. C., Li, X., Jin, M. W. and Li, G. R. (2012). Evidence for functional expression of TRPM7 channels in human atrial myocytes. Basic Res Cardiol 107, 282.

47. Henquin, J. C., Tamagawa, T., Nenquin, M. and Cogneau, M. (1983). Glucose modulates Mg2+ fluxes in pancreatic islet cells. Nature 301, 73–74.

48. Hellman, B., Idahl, L. A., Lernmark, A. and Taljedal, I. B. (1974). The pancreatic beta-cell recognition of insulin secretagogues: does cyclic AMP mediate the effect of glucose? Proc Natl Acad Sci U S A 71, 3405–3409.

49. Hill, R. S., Oberwetter, J. M. and Boyd, A. E., 3rd (1987). Increase in cAMP levels in beta-cell line potentiates insulin secretion without altering cytosolic free-calcium concentration. Diabetes 36, 440–446.

50. Tengholm, A. (2012). Cyclic AMP dynamics in the pancreatic beta-cell. Ups J Med Sci 117, 355–369.

51. Resnick, L. M., Altura, B. T., Gupta, R. K., Laragh, J. H., Alderman, M. H. and Altura, B. M. (1993). Intracellular and extracellular magnesium depletion in type 2 (non-insulin-dependent) diabetes mellitus. Diabetologia 36, 767–770.

52. Barbagallo, M., Gupta, R. K., Dominguez, L. J. and Resnick, L. M. (2000). Cellular ionic alterations with age: relation to hypertension and diabetes. J Am Geriatr Soc 48, 1111–1116.

53. Sharma, R. B. and Alonso, L. C. (2014). Lipotoxicity in the pancreatic beta cell: not just survival and function, but proliferation as well? Curr Diab Rep 14, 492.

54. Peng, Z., Aggarwal, R., Zeng, N., He, L., Stiles, E. X., Debebe, A., Chen, J., Chen, C. Y. and Stiles, B. L. (2020). AKT1 Regulates Endoplasmic Reticulum Stress and Mediates the Adaptive Response of Pancreatic beta Cells. Mol Cell Biol 40.

55. Szabat, M., Page, M. M., Panzhinskiy, E., Skovso, S., Mojibian, M., Fernandez-Tajes, J., Bruin, J. E., Bround, M. J., Lee, J. T., Xu, E. E., et al. (2016). Reduced Insulin Production Relieves Endoplasmic Reticulum Stress and Induces beta Cell Proliferation. Cell Metab 23, 179–193.

56. Boucher, J., Kleinridders, A. and Kahn, C. R. (2014). Insulin receptor signaling in normal and insulin-resistant states. Cold Spring Harb Perspect Biol 6.

57. Tuttle, R. L., Gill, N. S., Pugh, W., Lee, J. P., Koeberlein, B., Furth, E. E., Polonsky, K. S., Naji, A. and Birnbaum, M. J. (2001). Regulation of pancreatic beta-cell growth and survival by the serine/threonine protein kinase Akt1/PKBalpha. Nat Med 7, 1133–1137.

58. Bao, Z. Q., Jacobsen, D. M. and Young, M. A. (2011). Briefly bound to activate: transient binding of a second catalytic magnesium activates the structure and dynamics of CDK2 kinase for catalysis. Structure 19, 675–690.

59. Harper, J. W., Adami, G. R., Wei, N., Keyomarsi, K. and Elledge, S. J. (1993). The p21 Cdk-interacting protein Cip1 is a potent inhibitor of G1 cyclin-dependent kinases. Cell 75, 805–816.

60. Polyak, K., Lee, M. H., Erdjument-Bromage, H., Koff, A., Roberts, J. M., Tempst, P. and Massague, J. (1994). Cloning of p27Kip1, a cyclin-dependent kinase inhibitor and a potential mediator of extracellular antimitogenic signals. Cell 78, 59–66.

61. Rubin, H. (1975). Central role for magnesium in coordinate control of metabolism and growth in animal cells. Proc Natl Acad Sci U S A 72, 3551–3555.

62. Matsuda, M., Mandarino, L. and DeFronzo, R. A. (1999). Synergistic interaction of magnesium and vanadate on glucose metabolism in diabetic rats. Metabolism 48, 725–731.

63. Wimhurst, J. M. and Manchester, K. L. (1972). Comparison of ability of Mg and Mn to activate the key enzymes of glycolysis. FEBS Lett 27, 321–326.

64. Garfinkel, L. and Garfinkel, D. (1985). Magnesium regulation of the glycolytic pathway and the enzymes involved. Magnesium 4, 60–72.

65. Stanger, B. Z., Tanaka, A. J. and Melton, D. A. (2007). Organ size is limited by the number of embryonic progenitor cells in the pancreas but not the liver. Nature 445, 886–891.

66. Wang, S., Yan, J., Anderson, D. A., Xu, Y., Kanal, M. C., Cao, Z., Wright, C. V. and Gu, G. (2010). Neurog3 gene dosage regulates allocation of endocrine and exocrine cell fates in the developing mouse pancreas. Dev Biol 339, 26–37.

67. Castiglioni, S., Leidi, M., Carpanese, E. and Maier, J. A. (2013). Extracellular magnesium and in vitro cell differentiation: different behaviour of different cells. Magnes Res 26, 24–31.

68. Jin, J., Wu, L. J., Jun, J., Cheng, X., Xu, H., Andrews, N. C. and Clapham, D. E. (2012). The channel kinase, TRPM7, is required for early embryonic development. Proc Natl Acad Sci U S A 109, E225–233.

69. Jacobson, D. A., Kuznetsov, A., Lopez, J. P., Kash, S., Ammala, C. E. and Philipson, L. H. (2007). Kv2.1 ablation alters glucose-induced islet electrical activity, enhancing insulin secretion. Cell metabolism 6, 229–235.

70. Roe, M. W., Philipson, L. H., Frangakis, C. J., Kuznetsov, A., Mertz, R. J., Lancaster, M. E., Spencer, B., Worley, J. F., 3rd and Dukes, I. D. (1994). Defective glucose-dependent endoplasmic reticulum Ca2+ sequestration in diabetic mouse islets of Langerhans. The Journal of biological chemistry 269, 18279–18282.

71. Vierra, N. C., Dadi, P. K., Jeong, I., Dickerson, M., Powell, D. R. and Jacobson, D. A. (2015). Type 2 Diabetes-Associated K+ Channel TALK-1 Modulates beta-Cell Electrical Excitability, Second-Phase Insulin Secretion, and Glucose Homeostasis. Diabetes 64, 3818–3828.

72. Vierra, N. C., Dadi, P. K., Milian, S. C., Dickerson, M. T., Jordan, K. L., Gilon, P. and Jacobson, D. A. (2017). TALK-1 channels control beta cell endoplasmic reticulum Ca(2+) homeostasis. Sci Signal 10.

